# Modulation of huntingtin degradation by cAMP-dependent protein kinase A (PKA) phosphorylation of C-HEAT domain Ser2550

**DOI:** 10.1101/2022.05.04.490694

**Authors:** Yejin Lee, Hyeongju Kim, Douglas Barker, Ravi Vijayvargia, Ranjit Singh Atwal, Harrison Specht, Hasmik Keshishian, Steven A Carr, Ramee Lee, Seung Kwak, Kyung-gi Hyun, Jacob Loupe, Marcy E. MacDonald, Ji-Joon Song, Ihn Sik Seong

## Abstract

Huntington’s disease (HD) is a neurodegerative disorder caused by an inherited unstable *HTT* CAG repeat that expands further, thereby eliciting a disease process that may be initiated by polyglutamine-expanded huntingtin or a short polyglutamine-product. Phosphorylation of selected candidate residues is reported to mediate polyglutamine-fragment degradation and toxicity. Here to support the discovery of phospho-sites involved in the life-cycle of (full-length) huntingtin, we employed mass spectrometry-based phosphoproteomics to systematically identify sites in purified huntingtin and in the endogenous protein, by proteomic and phospho-proteomic analyses of members of an HD neuronal progenitor cell panel. Our results bring total huntingtin phospho-sites to 95, with more located in the N-HEAT domain relative to numbers in the Bridge and C-HEAT domains. Moreover, phosphorylation of C-HEAT Ser2550 by cAMP-dependent protein kinase (PKA), the top hit in kinase activity screens, was found to hasten huntingtin degradation, such that levels of the catalytic subunit (PRKACA) were inversely related to huntingtin levels. Taken together these findings highlight categories of phospho-sites that merit further study and provide a phospho-site kinase pair (pSer2550-PKA) with which to investigate the biological processes that regulate huntingtin degradation and thereby influence the steady state levels of huntingtin in HD cells.

## INTRODUCTION

Huntington’s disease (HD) (MIM 143100) is a dominantly inherited brain disorder, featuring characteristic neurodegeneration and motor, cognitive and behavioral clinical signs (McColgan & Tabrizi, 2018). The root genetic cause of HD is an expanded CAG triplet repeat in the *Huntingtin* gene (*HTT*) that extends a polyglutamine segment in huntingtin (“A novel gene containing a trinucleotide repeat that is expanded and unstable on Huntington’s disease chromosomes. The Huntington’s Disease Collaborative Research Group,” 1993), an HEAT (<u>H</u>untingtin, <u>E</u>longation factor 3, protein phosphatase 2A regulatory subunit PR65/<u>A</u> and target of rapamycin <u>T</u>OR1) repeat protein (Andrade & Bork, 1995). For individuals inheriting expansions in the fully penetrant range (40 or more repeats), the age at onset is hastened with increasing size of the expanded repeat (Hendricks et al., 2009; Lee et al., 2012).

Genetic studies with HD individuals support a pathogenic process leading to onset that involves further expansion of the inherited expanded repeat in brain cells over time until a critical threshold-length is reached, whereupon a process damaging to target neurons is initiated (Hong et al., 2021). The toxicity-mechanism(s) is not known but may involve an impact of the threshold-length repeat at the level of the mutant huntingtin protein or an aggregation-prone *HTT* exon1 encoded polyglutamine product, for example generated by exon 1-missplicing (Sathasivam et al., 2013).

Therapeutic strategies currently in trials aim to lower mutant huntingtin levels by *HTT*-silencing (Marxreiter, Stemick, & Kohl, 2020; Tabrizi, Ghosh, & Leavitt, 2019). However, understanding the role of posttranslational modification (PTM), particularly phosphorylation, in regulating huntingtin turnover (i.e. the balance between degradation and replacement synthesis), has also been proposed as route to lowering a toxicity-provoking entity (reviewed in (Lontay, Kiss, Virag, & Tar, 2020; Sambataro & Pennuto, 2017)). Investigations, based on predicated kinase sites within amino terminal (polyglutamine-containing) fragments of the protein, have identified several kinases, including IκB kinase (IKK)/TANK-binding kinase 1 (TBK1), Nemo-Like Kinase (NLK), and AKT/SGK kinases, phosphorylating serine residues S13/S16, S120 and S421, respectively, that modulate N-terminal polyglutamine-fragment degradation and toxicity (Hegde et al., 2020; Jiang et al., 2020; Kratter et al., 2016; Thompson et al., 2009). Phosphorylation status at those same IKK and NLK sites near the amino terminus is also implicated in modulating endogenous huntingtin levels (Jiang et al., 2020; Thompson et al., 2009). The advent of systems to express and purify (full-length) human huntingtin, with different polyglutamine segments (Huang et al., 2015; Kim, Hyun, Lloret, Seong, & Song, 2021; Vijayvargia et al., 2016), now facilitates the delineation of huntingtin’s HEAT repeat domain structure (Guo et al., 2018; Harding et al., 2021; T. Jung et al., 2020) and the discovery of residues that can be phosphorylated (phospho-sites) in the context of the entire protein (Huang et al., 2015; T. Jung et al., 2020; Ratovitski et al., 2017; Schilling et al., 2006). In addition, databases are also accruing phospho-sites identified on peptides derived from endogenous huntingtin detected in proteomic and phospho-proteomic studies of different cell and tissue types.

Here, building on previous studies with purified huntingtin, we have used unbiased approaches to delineate huntingtin phospho-sites, assessing the impact of polyglutamine length, thereby augmenting knowledge of this PTM in the context of the endogenous protein and making the unexpected discovery of a phospho-site/kinase pair that can modulate the steady-state level of huntingtin in growing cells.

## RESULTS

### Truncating huntingtin exposed N-HEAT domain phosphorylation sites

Structural analyses of purified human huntingtin/HAP-40 complex has delineated three main huntingtin structural domains, called N-HEAT, Bridge and C-HEAT (Guo et al., 2018) (Figure 1A), though the floppy ‘unstructured’ portions of the protein (∼25%) remain unresolved (S.Table 1). The majority of the 16 phospho-sites that we previously identified by mass spectrometry (LC-MS/MS) of huntingtin purified from our Baculovirus Sf9 insect cell expression system (T. Jung et al., 2020) are in the Bridge (5 sites) and C-HEAT (7 sites) domains, with relatively few in the large N-HEAT domain (4 sites) (Figure 1B), though the latter has many predicted kinase target sites. This distribution is consistent with Cryo-EM and biophysical analyses of purified huntingtin showing that the amino-terminal region is in close proximity with the C-HEAT region, such that the polyglutamine segment appears ‘buried’ (T. Jung et al., 2020). Consequently to discover N-HEAT sites we performed LC-MS/MS on three truncated human huntingtin products purified from Sf9 insect cell extracts. Seven of the eight phospho-sites identified with the ∼200 kDa product (C1213-3144) were a subset of sites reported for purified huntingtin (1-3144), most in unresolved regions in the Bridge and C-HEAT domains, and one new site (Ser1215-p) near the terminus of the product (Figure 1B, S.Table 1, S.Table 2). By contrast, across all polyglutamine lengths (23-, 46-, 78-residues), analyses of the ∼150 kDa product (N-1192) and smaller ∼60 kDa product (N-589) together disclosed 20 phospho-sites; 3 reported with purified huntingtin and 17 not detected in the context of the entire purified protein, nearly all in N-HEAT locations not resolved in the cryoEM huntingtin/HAP-40 structure (Figure 1B, S.Table 1, S.Table 2).

**Figure 1.**
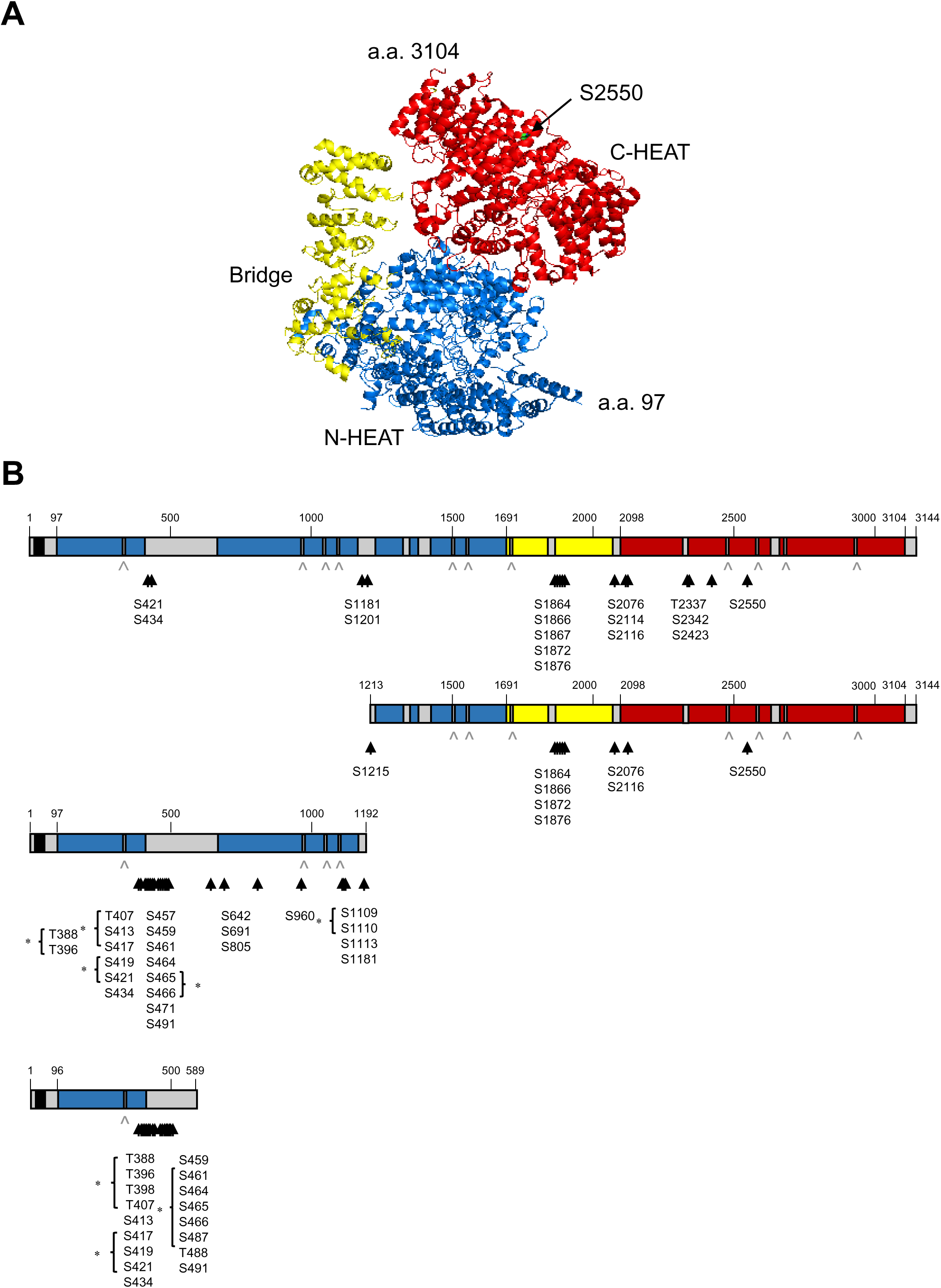
Location of purified huntingtin phospho-sites relative to cryo-EM structure domains. **A.** The human huntingtin cryo-EM structure without HAP 40, generated with PyMOL v1.7.4.5 Edu Enhanced for Mac OS X using RSCB using PDB: 6EZ8 (Guo et al., 2018), shows the N-HEAT, Bridge and C-HEAT domains, colored in blue, yellow and red, respectively. The amino acid (a.a.) start (a.a. 97) and end (a.a. 3104) of the resolved structure and position of C-HEAT Ser2550 (S2550) (green) are indicated. **B.** The top schematic depicts full-length huntingtin with its polyglutamine tract (black bar), N-HEAT (blue), Bridge (yellow) and C-HEAT (red) domains and unresolved regions >20 amino acids (grey bars) or ≤ 20 of amino acids (grey arrow heads) denoted. The amino acid numbering indicates the start (a.a.97) and end of resolved regions (a.a. 3104), the start of the Bridge (a.a.1691) and C-HEAT (a.a.2098) domains. The locations of phospho-sites we reported previously for purified Q23-huntingtin (T. Jung et al., 2020) are indicated under the schematic (black arrowheads). The schematic on the line below depicts the C1213-3144 fragment (a.a.1213-3144) and below that the schematics of the N1192 fragment (a.a.1-1192) (polyglutamine segments of 23, 46 and 78 residues) and the N589 fragment (a.a. 1-589) (23- and 46-polyglutamines), with phospho-sites identified in this study indicated (black arrow heads). Sites on a peptide that cannot be unambiguously identified (more than one potential residue) are marked by an asterisk.

Seven of the N-HEAT phospho-sites were not reported in a phospho-site aggregation-database (PhosphSitePlus (Hornbeck et al., 2015)) but 20 of the sites had been reported previously, as had many other N-HEAT residue phospho-sites, a number of which were detected in high-throughput proteomic (mainly cancer cell) studies. Since the latter implied an ‘openness’ of the N-HEAT domain in endogenous huntingtin, we set out to systematically discover endogenous huntingtin phospho-sites in neuronal progenitor cells (NPC), a cell type more relevant to HD.

### Proteomic NPC survey identified N-HEAT endogenous huntingtin phosphosites

We conducted an LC-MS/MS discovery survey, utilizing 12 members of an HD NPC panel with *HTT* CAG repeat alleles in the normal (<36 units) and expanded (>39 repeat) ranges. The strategy, designed with duplicates and cross-extract pooled reference controls (24 fractions in total) is depicted schematically in S.Figure 1 and described in the Methods section. Principle component analysis (PCA) revealed that variation in protein abundance levels (normalized to pooled reference control) (S.Figure 2) did not particularly distinguish NPC with *HTT* CAG repeats in the normal (non-HD) range (17-33 units) (hNPC.01 through hNPC.05) from those with repeats in the expanded HD-causing range (42-72 units) (hNPC 06 through NPC.11). One outlier NPC (hNPC.12), which grows slowly, having an extreme ∼180-220 CAG repeat, was excluded from subsequent analyses. As illustrated in S.Figure 3, the huntingtin-derived peptides detected in hNPC.01 through hNPC.11 (38.14% total huntingtin coverage) (S.Table 3) were located in structurally resolved and unresolved regions of the N-HEAT, Bridge and C-HEAT domains. The peptide abundance across the different NPC did not vary in an obviously systematic manner with CAG repeat length.

In contrast to the broad huntingtin peptide-coverage, the 26 huntingtin phospho-sites (22 phospho-serine and 4 phospho-threonine) identified across the eleven NPC (S.Table 3) are located (in the primary amino acid sequence) in clusters (Figure 2). The vast majority (21/26; 81%) lie within unresolved regions of the protein (S.Table 1), mainly in the N-HEAT domain (16/26), whereas few mapped to structured portions (N-HEAT four; C-HEAT one). Despite the anti-tyrosine antibody column enrichment step, no endogenous huntingtin phospho-tyrosine peptide sites were detected. The abundance of the identified phospho-site peptides relative to the reference control, was not evidently associated with *HTT* CAG size, implying that any impact of the length of the polyglutamine segment is subtle relative to the influence of other factors that determine the pattern of huntingtin phosphorylation in NPC.

**Figure 2.**
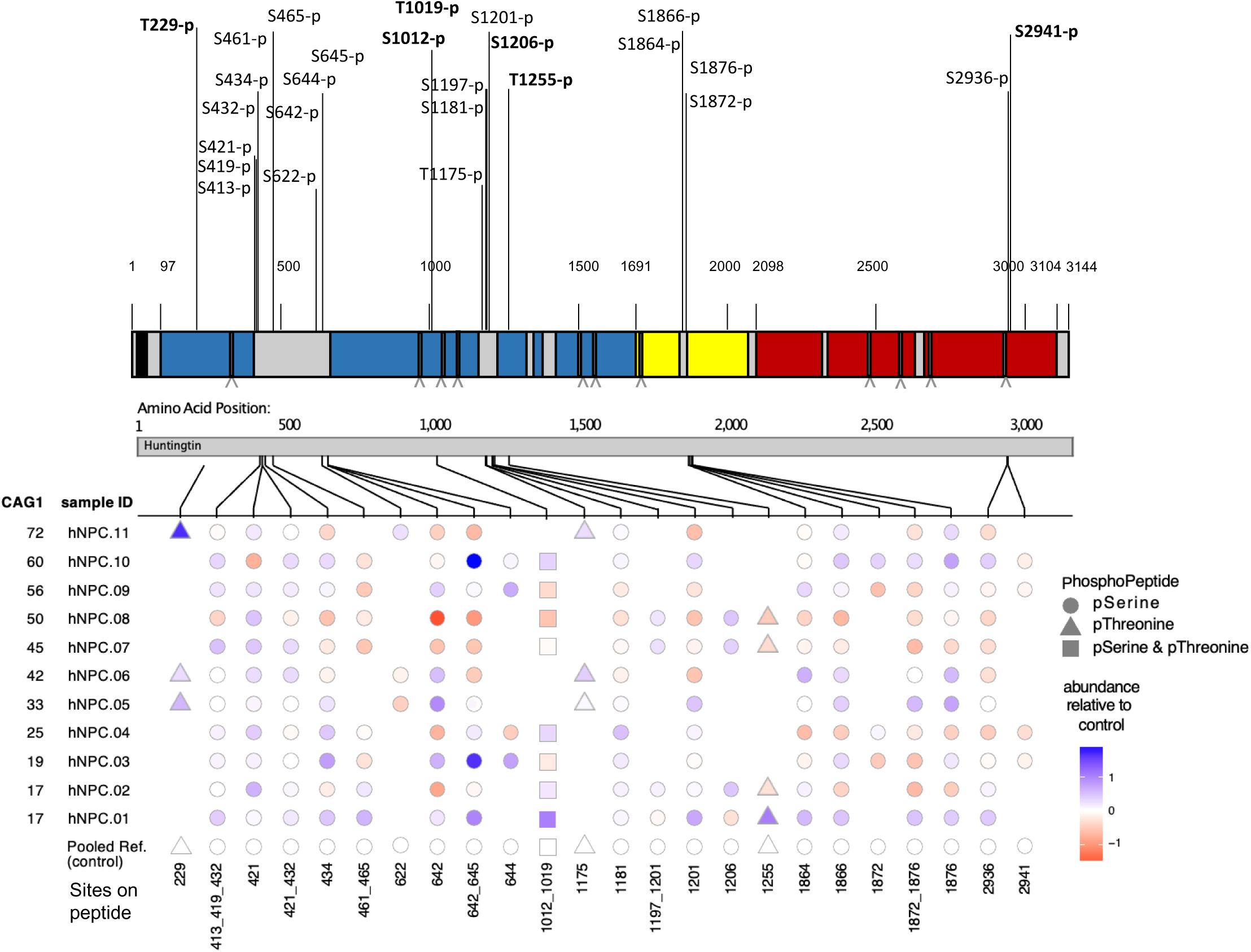
Location of phospho-sites NPC endogenous huntingtin across its domain structure. The schematic shows the residue location of the 26 phospho-sites identified in NPC, with those not previously reported indicated by bold text, along full-length huntingtin with its polyglutamine tract (black bar), N-HEAT (blue), Bridge (yellow) and C-HEAT (red) domains (unresolved regions >20 amino acids = grey bars or ≤ 20 of amino acids = grey arrow heads beneath the schematic), numbering as in Figure 1B. Below that, aligned to the greyed huntingtin schematic, is a plot summarizing the locations and scaled relative abundance (compared to pooled reference control) of the phospho-serine (circles), phospho-threonine (triangles) and phospho-serine-threonine (squares) peptides identified for each NPC (hNPC.01 through hNPC.11) ranked (Y-axis) by decreasing size of the longer *HTT* CAG repeat allele, with the identified phospho-site peptide plotted on the X axis.

Comparison of these 26 phospho-sites with the previously identified endogenous huntingtin phospho-sites reported in proteomic studies (https://www.phosphosite.org/proteinAction.action?id=1292&showAllSites=true (Hornbeck et al., 2015)), when mapped onto the huntingtin/Hap40 domain structure coordinates (S.Table 1; S.Table 4), revealed a total of 67 endogenous huntingtin phospho-sites: 43 N-HEAT (35 reported plus 8 new from this study), 9 Bridge (4 this study all reported previously) and 15 C-HEAT (14 reported plus 1 new from this study). The distribution underscored the relative richness of phospho-sites in the N-HEAT domain, which notably included phospho-sites detected with short N-terminal exon1 huntingtin fragment (Aiken et al., 2009; Chiki et al., 2021; Hegde et al., 2020) (S.Table 4) and the comparative paucity of phospho-sites in the large C-HEAT domain.

### *In vitro* screens identified PKA as the top kinase phosphorylating purified huntingtin

In an attempt to discover additional huntingtin phospho-sites, in a manner that may imply functional consequences, we conducted plate format screens of a panel of protein kinases (245 Serine/Threonine; 94 Tyrosine) (Reactive Biology). Independent screens utilized either FLAG-tagged Q23-huntingtin or Q78-huntingtin purified from Sf9 Baculovirus system (see the detail in Methods) as substrates. We sought kinases that incorporated ^33^Pi (from gamma-^33^P-ATP) above levels observed in replica plates without purified huntingtin (^33^Pi-incorporation ratio). Amongst 73 kinases, with a cut-off ^33^Pi-incorporation ratio >5, (S. Table 5), a few potentially preferentially phosphorylated either Q23-huntingtin, including G-protein-coupled receptor kinase 6 (GRK6), and MAP/microtubule-affinity regulating kinase 3 (MARK3), or Q78-huntingtin, including G-protein-coupled receptor kinase 3 (GRK3), casein kinase 1 isoform epsilon (CK1-epsilon), and dual-specificity tyrosine-regulated kinase 2 (DYRK2) (Figure 3). However, the vast majority exhibited similar activity on both substrates. Serum glucocorticoid regulated kinase 2 (SGK2) and MAP/microtubule-affinity regulating kinase 2 (MARK2), with ^33^Pi-incorporation ratios >40, were both notable but cAMP-dependent protein kinase A (PKA) stood apart because it exhibited four-fold higher activity than these on both substrates (^33^Pi-incorporation ratio >180) (Figure 3).

**Figure 3.**
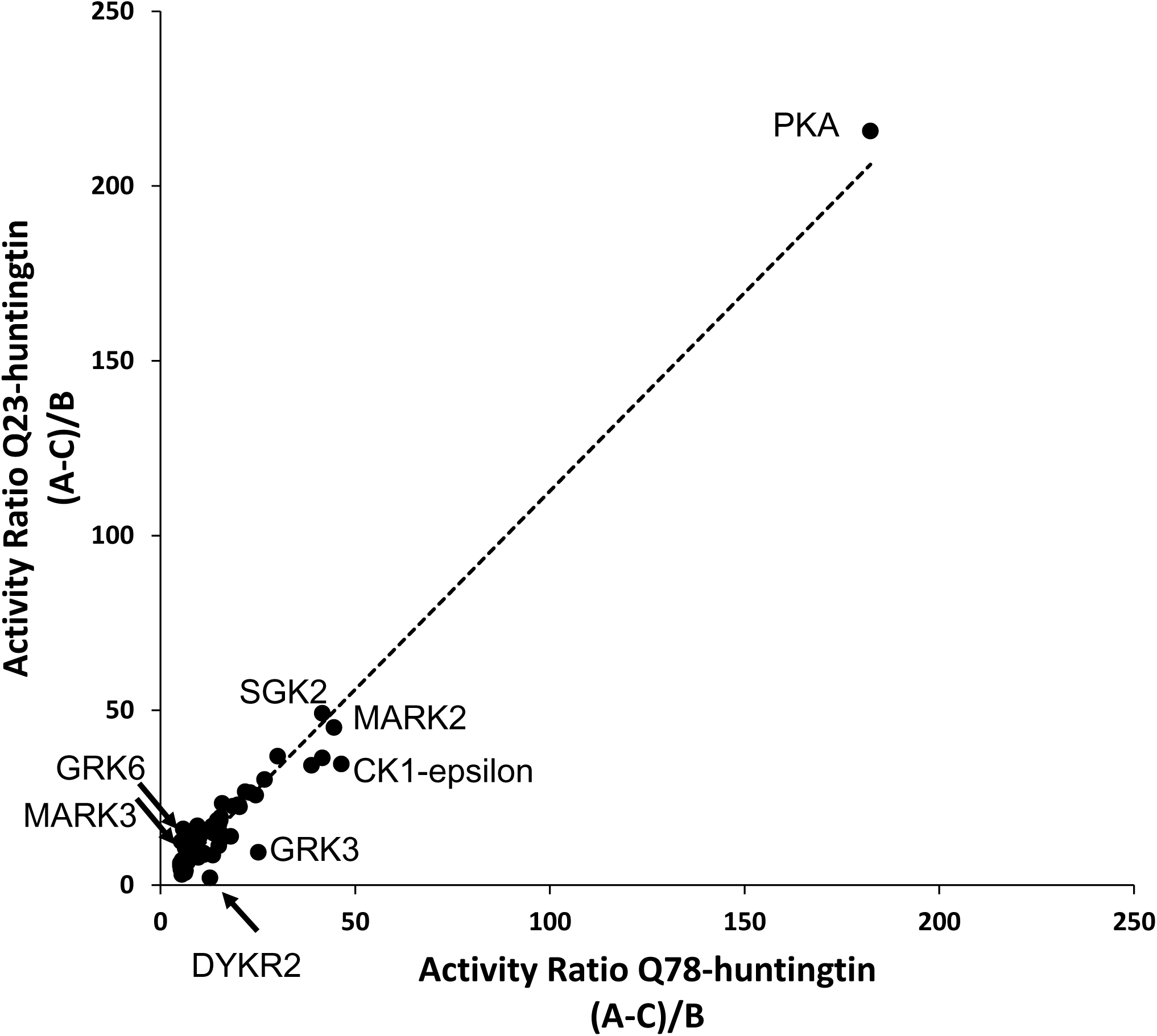
PKA is top hit from Q23-huntingtin and Q78-huntingtin i*n vitro* kinase screens. The plot compares kinase activity ratio for ^33^Pi incorporation into Q23-huntingtin compared to Q78-huntingtin, for the 73 kinases showing activity ratio >5 on both Q23- and Q78 huntingtin. The activity ratio was calculated by the equation, *(A-C)/B*, where *A* = Intensity raw value, *B* = Kinase autophosphorylation value and *C* = Substrate-background value for normalized mean of triplicates from mean of two individual experiments. A few kinases showed potential preference for Q23-huntingtin (e.g. GRK6, MARK3) or Q78-huntingtin (e.g. DYKR2, GRK3, CK1-epsilon) but most had similar activities on both substrates, including SGK2 and MARK2, with notable level of activity, though PKA was the top hit.

### The PKA catalytic subunit phosphorylated huntingtin at serine 2550

To pursue this striking observation, we first demonstrated PKA catalytic subunit (PRKACA)-mediated incorporation of ^32^Pi (from gamma-^32^P-ATP) into Q23-huntingtin (S.Figure 4A), and then utilized LC-MS/MS analysis of purified Q23-huntingtin incubated with or without PKA catalytic subunit PRKACA, to identify potential PKA target sites. PRKACA increased the detection of two of the four highly reliable phosphopeptides (Ascore >19; counts >2); N-HEAT Ser1201 and C-HEAT Ser2550 (Figure 4A), though the former phosphopeptide was also detected in the absence of PRKACA. Since the Ser2550 phosphopeptide was detected only in the presence of the PKA catalytic subunit (Figure 4A) and also is flanked by a canonical PKA motif (RKLS), conserved in huntingtins of other vertebrates (Figure 4B), we focused subsequent analysis on this C-HEAT site.

**Figure 4.**
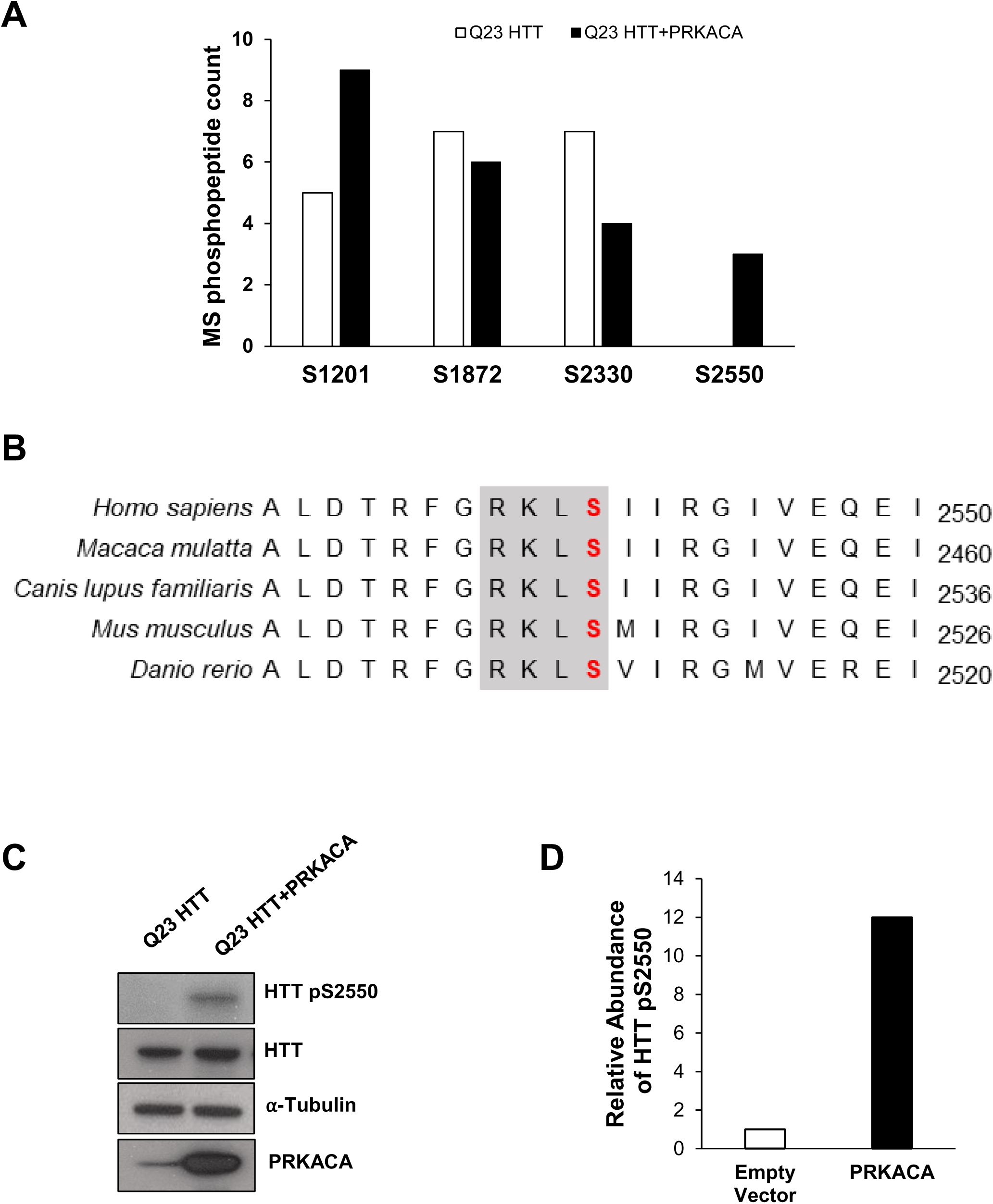
PKA phosphorylates huntingtin at Ser2550. **A.** Bar graph showing MS/MS results of each phosphopeptide from Q23-huntingtin (HTT) incubated without (open bar) or with purified PRKACA (black bar). **B.** The multiple sequence alignment compares the human huntingtin amino acid sequence focused on the region of Ser2550 (Red bold) with huntingtins predicted from the gene sequence of other orthologs*, Macaca mulatta, Canis lupus familiaris, Mus musculus, and Danio rerio*. Shading denotes the conservation of PKA substrate motif (RKXS) across these organisms. **C.** Immunoblot showing result of detecting pSer2550 in FLAG tagged huntingtin overexpressed without or with PRKACA overexpression in *HTT* null HEK 293T cells for 24 hours, with abHTT-pS2550, as well as huntingtin level detected with anti-huntingtin reagent (MAB2166), and PRKACA level detected with anti-PRKACA reagent, and α-tubulin as loading control with anti-α-tubulin reagent. **D.** Plot showing relative abundance of pSer2550 endogenous huntingtin peptide detected by PRM assay with or without overexpression of PRKACA for 24 hours in HEK 293T cells. Endogenous huntingtin was enriched by immunoprecipitation with MAB2166 antibody before PRM assay.

PRKACA phosphorylation of Ser2550 in the context of Ser2550-peptide (S. Figure 4B) and purified Q23-huntingtin and Q78-huntingtin (S. Figure 4C) was confirmed by anti-pS2550 immunoblot analyses. Moreover, when PRKACA was co-transfected into *HTT*-null HEK293T cells with exogenous Q23-huntingtin, immunoblot analysis with pS2550 antibody (24 hours post transfection) detected a robust band of Q23-huntingtin-pSer2550 (Figure 4C), while anti-huntingtin reagents detected Q23-huntingtin, though by 48 hours the band intensity was decreased (S.Figure 4D). Indeed, detection of pSer2550-endogenous huntingtin, which confirmed that this C-HEAT site is phosphorylated by PKA, was difficult by anti-pS2550 immunoblot analysis (data not shown) and required both exogenous PRKACA expression in HEK293T cells and a sensitive huntingtin Ser2550 residue parallel-reaction monitoring (PRM) assay performed with pSer2550-immuno-enriched protein (Figure 4D).

### PRKACA phosphorylation of Ser2550 hastened huntingtin degradation

The evidence for PKA mediated Ser2550 phosphorylation seemed at odds with the apparent inability to detect pSer2550 in the absence of PRKACA co-expression, in either exogenously expressed huntingtin (Figure 4C) or in endogenous huntingtin (Figure 2, S.Table 4, Figure 4D). A possible resolution to this apparent conundrum, implied by the decrease in Q23-huntingtin levels by 48 hours post PRKACA transfection (S.Figure 4D), was that phosphorylation of Ser2550 by this kinase may influence huntingtin turnover by altering the rate at which the protein is degraded. Assessing this presumption, we first confirmed by anti-huntingtin immunoblot analysis that Ser2550 was required for PRKACA co-expression to decrease Q23-huntingtin level, which revealed that this effect was not observed with 2550-alanine mutated Q23-huntingtin-2550A (Figure 5A). Then immunoblot monitoring of Q23-huntingtin and Q78-huntingtin levels in *HTT*-null HEK293T cells, after timed treatment with the protein translational-blocker cycloheximide, starting 24 hours post-transfection (P0), clearly demonstrated that PRKACA co-expression hastened the rate of degradation of Q23-huntingtin and Q78-huntingtin. Levels of Q23-huntingtin (Figure 5B) and Q78-huntingtin (Figure 5C) decreased rapidly in cells expressing exogenous PRKACA, with a drop in normalized band-intensity evident at 2 hours post-cycloheximide and declining further over the next six hours at a rate estimated to be 0.07 units/hour and 0.1 units/hour, respectively, compared to 0.02 units per hour and 0.05 units per hour (respectively) for the same time-interval in control (empty-vector) transfected cells. Therefore, blocking the synthesis of new protein revealed that Ser2550 phosphorylation by exogenous PRKACA dramatically increased (three to four fold) the rate of Q23-huntingtin and Q78-huntingtin degradation, which seems likely to explain the impact of PRKACA in decreasing huntingtin levels over time in the absence of cycloheximide (S.Figure 4D and Figure 5A).

**Figure 5.**
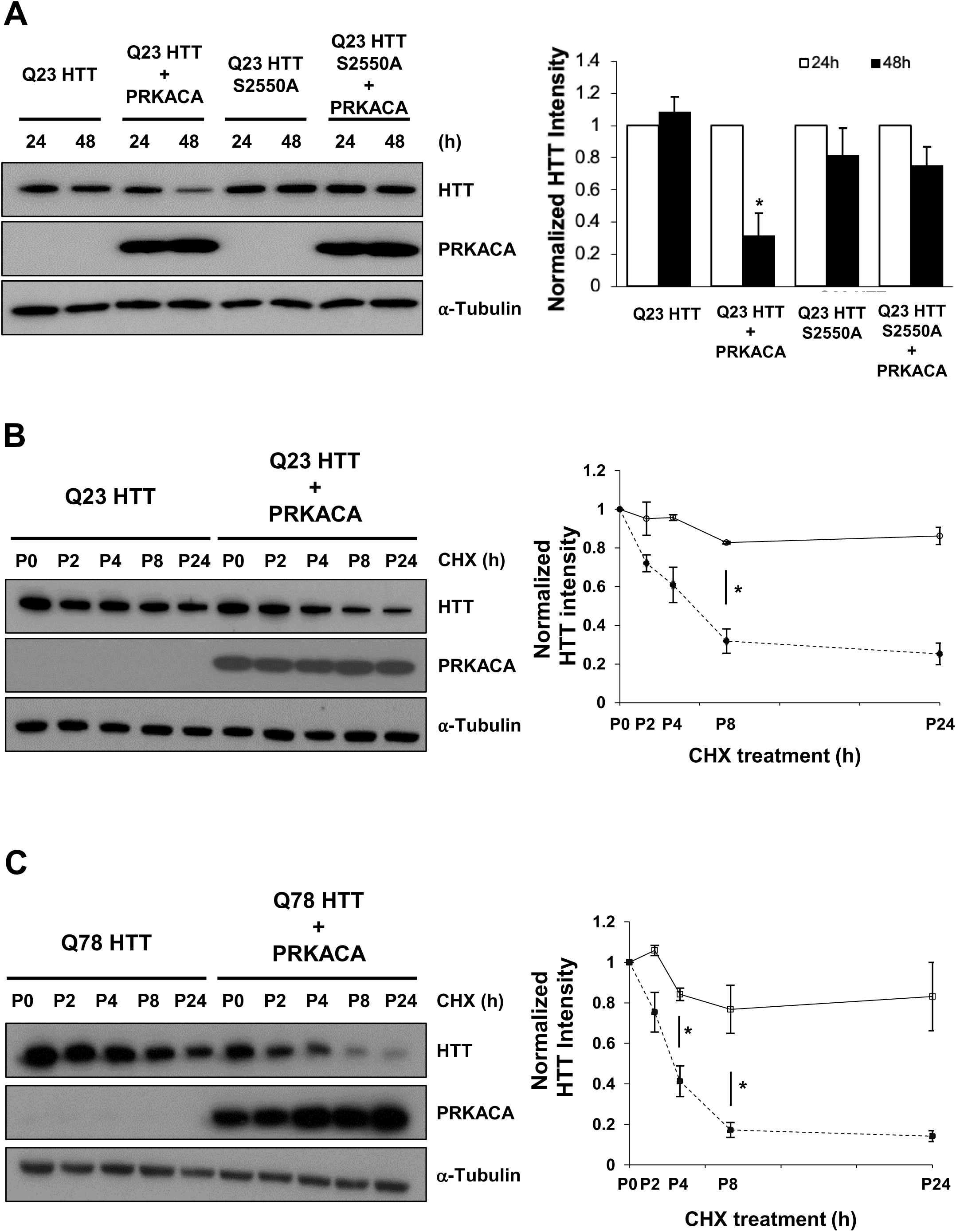
PKA Ser2550 phosphorylation increases the rate of huntingtin degradation. **A.** The immunoblot (left) shows the results of detecting Q23-huntingtin (HTT) or Q23-huntingtin phosphomutant S2550A overexpressed in *HTT* null HEK 293T cells in the absence or presence of PRKACA for 24 h or 48 h, with anti-huntingtin reagent (MAB2166) and PRKACA level detected with anti-PRKACA reagent, and α-tubulin as loading control with anti-α-tubulin reagent. The histogram (right) shows the huntingtin/⍺-tubulin band intensities normalized to the mean ratio of each sample at post 24 hours transfection. Data represent mean ± s.e.m (n = 2). **B and C.** immunoblots (left) showing cycloheximide (CHX) chase experiments of normal Q23-huntingtin (Q23 HTT) (**B**) and mutant Q78-huntingtin (Q78 HTT) (**C**) overexpressed without or with PRKACA overexpression in *HTT* null HEK 293T cells, probed with anti-huntingtin reagent (MAB2166), anti-PRKACA reagent and anti-α-tubulin reagent for huntingtin, PRKACA and α-tubulin as loading control, respectively. For each, the plots (right) show the huntingtin/⍺-tubulin band intensities normalized to the mean ratio of each sample at post 24 hours transfection (P0) in the absence (open symbol) and presence (closed symbol) of PRKACA overexpression). Data represent mean ± s.e.m (n = 2). Asterisks indicate level of statistical significance (paired Student’s t-test, two tailed); ∗ P < 0.05.

### Reciprocal relationship between level of PRKACA and level of huntingtin

Consequently, because our interest is in modulators of endogenous huntingtin, we investigated whether, as implied by the results above, exogenously expressed PRKACA would decrease endogenous huntingtin protein and, exploring a potential relationship, we also assessed whether reducing expression of endogenous PRKACA (using specific shRNA) would impact huntingtin level. As shown in Figure 6, immunoblot analysis revealed that HEK293T cells with exogenous-PRKACA exhibited lower levels of endogenous huntingtin (48 hours posttransfection), than untreated HEK293T cells (Figure 6A), whereas HEK293T cells stably expressing *PRKACA*-specific-shRNA expression vector, but not HEK293T cells with the scramble-shRNA vector, was associated with increased huntingtin band intensity, thereby implying an increase in huntingtin level (Figure 6B).

**Figure 6.**
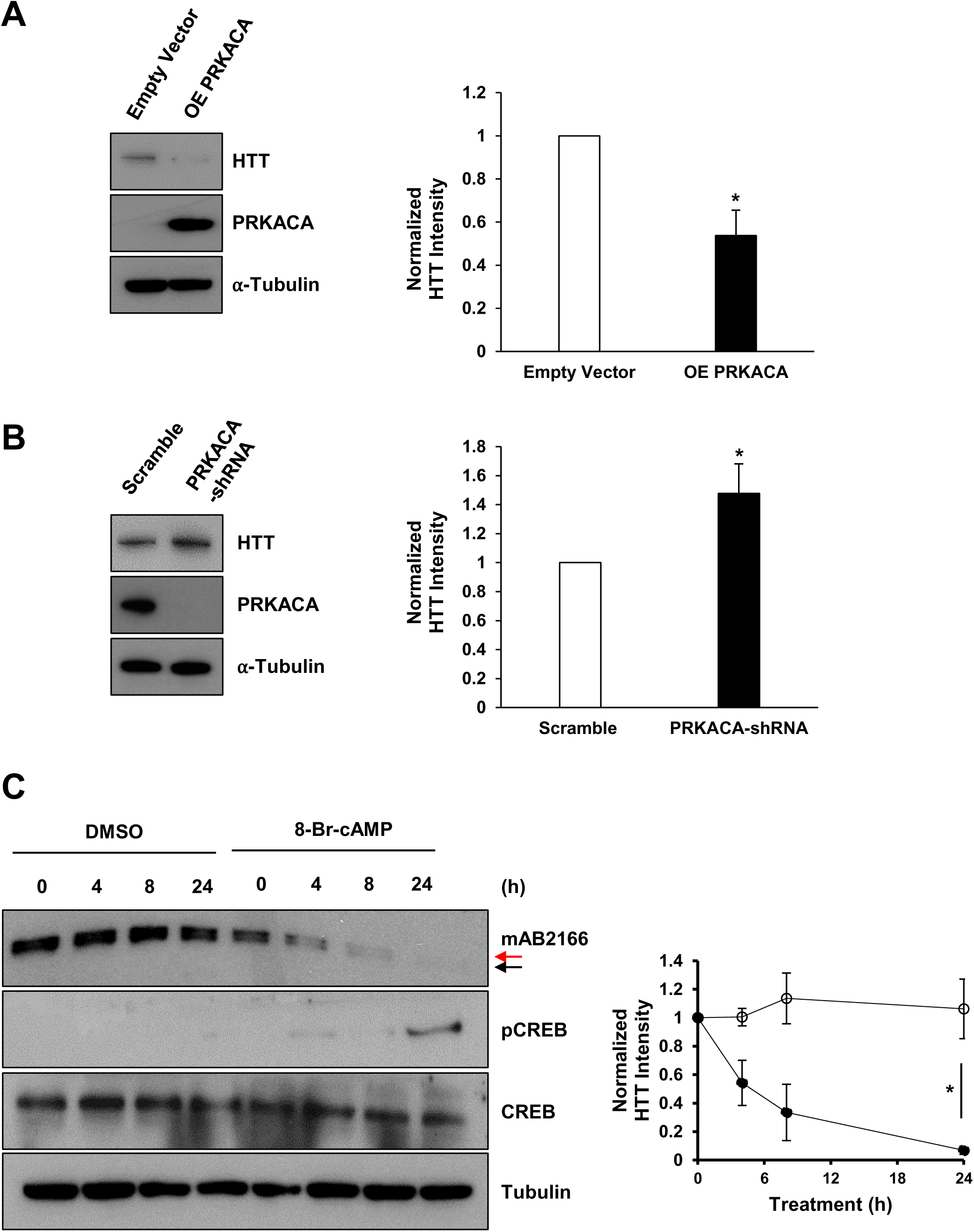
Cellular PKA activity is associated with endogenous huntingtin level. **A.** The immunoblot (left) shows the expression levels of endogenous huntingtin (HTT) without or with overexpressed PRKACA in HEK293 cells, detected with anti-huntingtin reagent (MAB2166) and PRKACA levels detected with anti-PRKACA reagent and α-tubulin as loading control with anti-α-tubulin reagent. The histogram (right) shows the huntingtin/⍺-tubulin band intensities normalized to the mean ratio of each sample of empty vector for empty vector (open bar) and PRKACA (closed bar) expressed samples. Data represent mean ± s.e.m (n = 3). **B.** The immunoblot (left) shows the expression level of endogenous huntingtin in HEK 293T cells stably expressing scramble-or PRKACA-shRNAs, detected with anti-huntingtin reagent (MAB2166) and PRKACA levels detected with anti-PRKACA reagent and α-tubulin as loading control with anti-α-tubulin reagent. The histogram (right) shows the huntingtin/⍺-tubulin band intensities normalized to the mean ratio of each sample expressing scramble-shRNA for scramble-shRNA (open bar) and PRKACA-shRNA (closed bar) expressed samples. Data represent mean ± s.e.m (n = 3). **C.** The immunoblot shows the expression level of endogenous huntingtin in hNPC (Q62/Q20) treated with DMSO or 300 µM of 8-Br-cAMP, detected with anti-huntingtin reagent (MAB2166) and phosphor-CREB with anti-phosphor-CREB (S133) reagent, total CREB with anti-CREB reagent and α-tubulin as loading control with anti-α-tubulin reagent. Of note, Q62-huntingtin (red arrow) separated from Q20-huntingtin (black arrow) was also confirmed by probing with 1F8 antibody (data not shown (White et al., 1997)). A histogram showing total huntingtin/⍺-tubulin band intensities normalized to the mean ratio of each sample at 0 hours for DMSO (open symbol) and 8-Br-cAMP (closed symbol) treated samples, respectively. Data represent mean ± s.e.m (n = 2). Asterisks indicate level of statistical significance (paired Student’s t-test, two tailed); ∗ P < 0.05.

To investigate whether these findings may be relevant to HD cells, we first determined whether PKA-activation would impact endogenous huntingtins with normal- and HD mutant expanded-length polyglutamine tracts, in a member of our HD NPC series (hNPC.10). Immunoblot analysis demonstrated that enhancing PKA activity by treatment of hNPC.10 cells with PKA activator 8-Bromoadenosine 3’,5’-cyclic adenosine monophosphate (8-Br-cAMP), which over time increased phosphorylation of PKA target protein CREB (24 hours), was rapidly (by 4 hours) associated with decreasing levels of both huntingtin and mutant huntingtin, expressed from the *HTT* normal (CAG 18) and expanded (CAG 60) alleles, respectively (Figure 6C).

The results of acute (genetic- and compound-mediated) manipulation of PKA activity predicted a (inverse) relationship between the physiological levels of PRKACA and (total) huntingtin, which vary across the 11 members of our NPC panel. We therefore utilized isobaric mass tag-based quantitative proteome analysis to determine the abundance levels of PRKACA and the levels of (total) huntingtin expressed from both alleles (*HTT* CAG repeats ranging from 17 to 72 units). The PRKACA abundance level was not significantly related to *HTT* CAG repeat size (Multiple R-squared: 0.2013 and p-value: 0.1663). However, as shown in the plot in Figure 7, the PRKACA abundance exhibited an inverse relationship with (total) huntingtin protein level (Multiple R-squared: 0.443 and p-value: 0.02539), such that as PRKACA increased, huntingtin was decreased.

**Figure 7.**
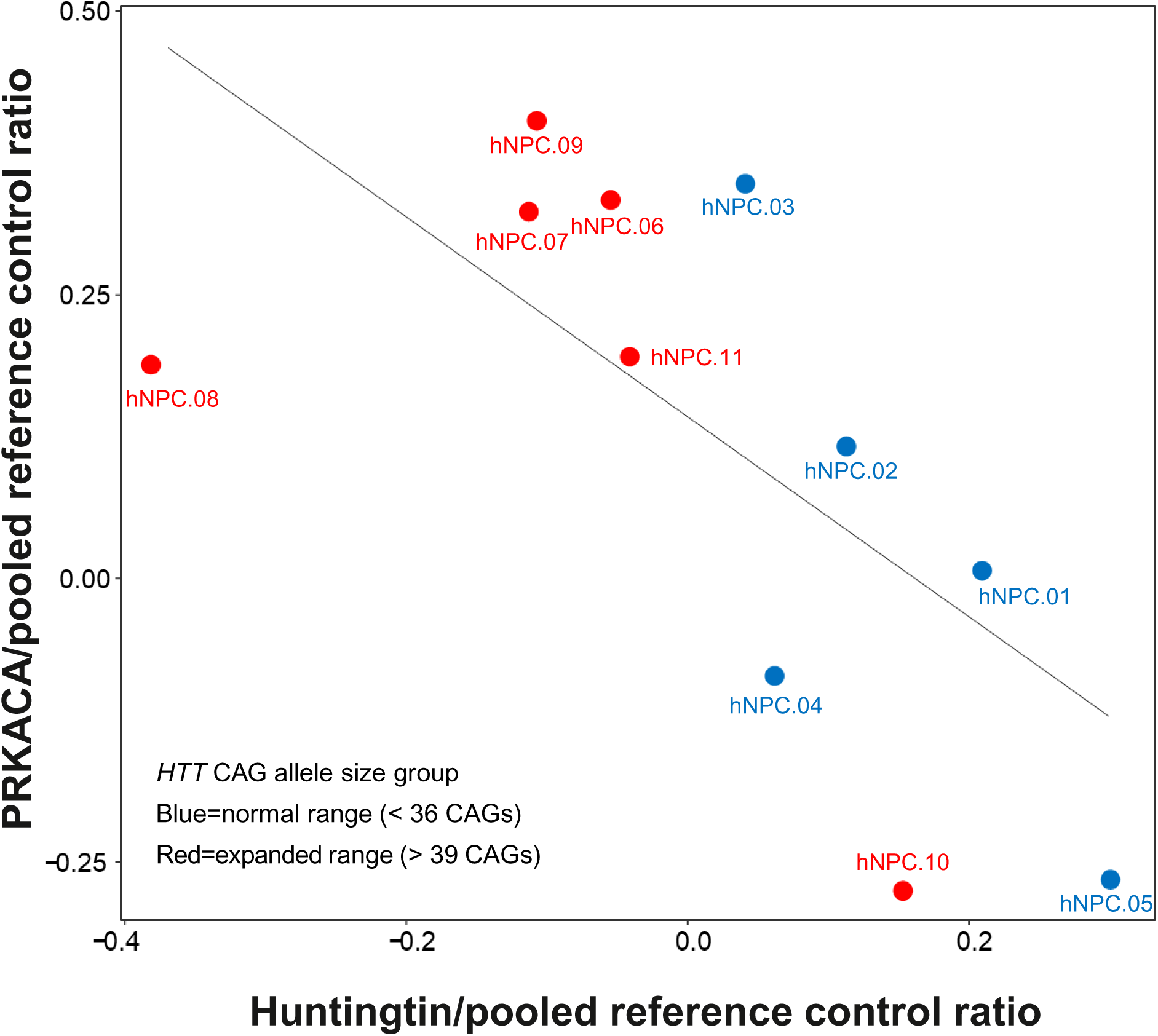
Inverse correlation between levels of huntingtin and PRKACA in a panel of HD neural progenitor cells. The scatter plot of the endogenous huntingtin ratio (normalized to pooled reference sample) (X axis) versus endogenous PRKACA ratio (normalized to pooled reference sample) (Y axis) measured by quantitative LC-MS/MS using TMT 10-plex across each of the 11 members of an hNPC panel shows an inverse correlation between huntingtin and PRKACA levels, as indicated by the linear regression line (black line), that is significant (Multiple R-squared: 0.443, Adjusted R-squared: 0.3812; F-statistic: 7.159 on 1 and 9 DF, p-value: 0.02539.)

## DISCUSSION

Genetic modifier studies point to the rate of (further) expansion of the pure CAG repeat in target cells as the driver of the timing of HD onset (Genetic Modifiers of Huntington’s Disease, 2015; Genetic Modifiers of Huntington’s Disease Consortium. Electronic address & Genetic Modifiers of Huntington’s Disease, 2019). The subsequent event that initiates toxicity in target cells may involve (further) polyglutamine-expanded huntingtin or an *HTT* exon1 polyglutamine-containing product (Michalik & Van Broeckhoven, 2003; Tabrizi, Flower, Ross, & Wild, 2020). The latter has been studied extensively, including predicted phospho-sites and kinases that can influence fragment aggregation, degradation or toxicity (Aiken et al., 2009; Gu et al., 2009; Hegde et al., 2020; Thompson et al., 2009). We, and others, are delineating phospho-sites that may be critical to the function and life-cycle of endogenous (full-length) huntingtin (Huang et al., 2015; T. Jung et al., 2020; Ratovitski et al., 2017; Schilling et al., 2006). Inclusion of MS-based phosphoproteomics analyses presented here, which confirmed 31 reported sites and added 14 new sites, a total of 95 phospho-sites have now been identified, 67 of which can be detected with endogenous protein. In conjunction with the cryoEM structure of huntingtin (Guo et al., 2018), this compendium provides insights into the clustered distribution of phospho-sites across the protein’s HEAT domains. Moreover, comparison across the various huntingtin-substrates (full-length/fragment; purified/endogenous) highlights several phospho-site categories with which to prioritize specific sites for functional study, in addition to pSer2550 which we have demonstrated to be involved in huntingtin turnover (i.e. modulating the balance between degradation and renewal synthesis).

The major category, perhaps important for ‘generic’ huntingtin function, comprises sites that can be phosphorylated in many cell-types/tissues. Such sites are detected with purified huntingtin and are also frequently identified in (largely cancer-oriented) phospho-proteomic studies (>12 studies). These include two phospho-site clusters in different structurally unresolved regions within the N-HEAT domain: (pSer419, pSer421, pSer434, and pSer1181, pSer1201) and a cluster in an unresolved region of the Bridge domain (pSer1864, pSer1872 and pSer1876). Indeed, because they can be detected in fragment, many of the N-HEAT sites are well-studied. Ser421 phosphorylation by AKT, and dephosphorylation by calcinurin, recruits and releases kinesin, respectively, thereby determining the direction of vesicle transport (e.g. BDNF) (Colin et al., 2008; Scaramuzzino, Cuoc, Pla, Humbert, & Saudou, 2022), while phosphorylation of this site is also reported to decrease huntingtin-cleavage and fragment-toxicity (Warby et al., 2005) and to influence mitochondrial phenotypes and toxicity in HD neuronal cells (Xu et al., 2020). CDK5 phosphorylation of pSer434 was associated with decreased caspase cleavage of huntingtin (Luo, Vacher, Davies, & Rubinsztein, 2005), whereas CDK5 phosphorylation of pSer1181 and pSer1201was reported to mediate huntingtin toxicity in neuronal cultures (Anne, Saudou, & Humbert, 2007). The cluster of Bridge sites is less well studied but in the context of highly purified protein, phosphorylation pSer1864 and pSer1876 varied with polyglutamine length, in a coordinataed manner with other sites, which uncovered a subtle impact of polyglutamine size on accessibility of Bridge domain phospho-sites and cross-talk with N-HEAT phospho-sites (T. Jung et al., 2020).

A second category, perhaps connoting a role in cell-type or cell-state specific huntingtin function, comprises sites identified in endogenous huntingtin in several phospho-proteomic studies but not detected in multiple studies with purified human huntingtin protein (full-length and fragment). pSer622 in the N-HEAT domain (unresolved region) and pSer2936, located in the C-HEAT (structured region) domain were both identified in a study of mitotic regulators, aurora and polo-like kinases (Kettenbach et al., 2011), while the latter was also detected in studies of stress kinase pathways (ischemic tumors) (Mertins et al., 2014), and JAK3 inhibitors and MEK/BCL2 inhibitors (T-cell acute lymphoblastic leukemia) (Degryse et al., 2018), with an Huntington’s Disease signaling module. These sites merit focused study, as huntingtin is reported to function in oxidative stress (Godin, Poizat, Hickey, Maschat, & Humbert, 2010; Machiela et al., 2020; Molina-Calavita et al., 2014).

A third category, of interest for huntingtin-lowering, comprises sites with potential function in huntingtin turnover; phospho-sites identified with highly purified huntingtin protein but not detected in endogenous huntingtin in any of the many mass spectrometry studies. Two C-HEAT domain sites, pSer2550 (structured region, Figure 1A) and pSer2076 (unresolved region), meet these criteria. The former site, as we discovered from the results of unbiased kinase screens with purified huntingtins, is a target of serine/threonine kinase PKA. PKA-mediated Ser2550 phosphorylation is involved in huntingtin turnover, as discussed below. However, pSer2076 has not been investigated in this light, though it has been reported of high interest for HD, and is also deserving of focused study, because it is polymorphic in the population due to a single nucleotide DNA polymorphism (∼0.1% minor allele frequency in HAPMAP) that changes the serine to a proline residue (Martin et al., 2018).

The discovery that PKA-mediated Ser2550 phosphorylation, by dramatically hastening the rate of huntingtin degradation, can influence huntingtin turnover and the steady state level of both normal-and expanded-polyglutamine huntingtin in growing cells is significant for several reasons. First, it hints at a dynamic (function-related) life-cycle, that is belied by the long steady-state half-life of the endogenous protein (Persichetti et al., 1996). In HD fibroblasts, the half-life of human huntingtin with normal- and expanded-range polyglutamine tracts was estimated to be ∼48 and ∼27 hours, respectively, with an half-life of total huntingtin protein of about 57 hours (Wu et al., 2016). Secondly, and supported by the finding that physiological variation in PRKACA level may be a meaningful determinant of huntingtin level, this discovery provides a specific kinase and phospho-site pair with which to delineate the molecular and cellular events that regulate and participate in huntingtin degradation and experiments to determine whether these are the same or different in non-dividing cells, particularly neuronal cells that are the vulnerable targets of the HD CAG repeat expansion mutation.

## MATERIALS AND METHODS

### Human huntingtin amino acid numbering

The polyglutamine tract in human huntingtin is polymorphic so that different reference amino acid sequences have different numbering. The cryo-EM huntingtin:HAP40 (Guo et al., 2018) was performed and numbered for a 17-polyglutamine tract; the PhosphoPlus site (Hornbeck et al., 2015) uses 21-polyglutamine tract. Here all human huntingtin amino acid numbering throughout is relative to NP 002102.4 reference sequence, with a 23-polyglutamine tract (T. Jung et al., 2020; Vijayvargia et al., 2016).

### Human FLAG-huntingtin cDNA clones in pFASTBAC1

The information about full-length *HTT* cDNA cloned into pFASTBAC1 vector (Invitrogen) was mentioned in the previous paper (Vijayvargia et al., 2016). Using them, *HTT* cDNA for N-1192 (amino acid 1-1192), N-589 fragments (amino acid 1-589) and C1213-3144 (amino acid 1213-3144) fragments were cloned into pFASTBAC1 as below: To make both large and small N-terminal fragments, the NcoI-XhoI HTT cDNAs with varying polyglutamine tracts (Q23, 46, 78) in pFASTBAC1were used (Vijayvargia et al., 2016). The 3 kb PCR product generated with two primers, forward: 5’-TTACAG<u>CTCGAG</u>(XhoI)CTCTATAAGG-3’ and reverse: 5’-ATAT<u>CCGCGG</u>(SacII)TTATGGTTCTTTCTCCTTCCC-3’ was inserted in frame using XhoI-SacII into the vector containing NcoI-XhoI HTT cDNA to make N-1192 fragment of huntingtin. The XhoI-KpnI *HTT* cDNA fragment, encoding huntingtin amino acid 170-589, was inserted into the same vector to generate N-589 fragment of huntingtin. C1213-3144 fragment in pFASTBAC1 was made through two steps, 1) 2.5 kb PCR product was generated with two primers, forward: 5’-ATGC<u>CTCGAG</u>(XhoI)AGACAATCTGATACC-3’ and reverse: 5’-ATG<u>CCGCGG</u>(SacII)AGCAAGGAT<u>GTCGAC</u>(SalI)CAT-3’ and inserted into pFASTBAC1 using XhoI and SacII. 2) the 3,528 bp SalI-SacII *HTT* cDNA fragment from a full *HTT* cDNA, pBS-HD1-3144Q23 (Seong et al., 2010) encoding huntingtin amino acid 2010-3144, was inserted in frame using SalI-SacII into the pFASTBAC1 containing the 2.5 kb PCR product above. All final clones were verified using full DNA sequence analysis. By convention, the amino acid numbering throughout the text follows the numbering of Q23-huntingtin (NP_002102.4) regardless of the length of the polyglutamine tract.

### Mass spectrometry identification of phosphorylation sites

For the determination of phosphorylation sites, recombinant N-1192 and N-589 fragments with polyglutamine tract length of 23, 46 and 78 and C1213-3144 fragment were purified in the presence of complete protease and phosphatase inhibitor cocktails (Roche Applied Science) to retain phosphorylation as described previously for recombinant full-length human huntingtin proteins (Vijayvargia et al., 2016). 10-20 μg of purified proteins were separated by SDS-PAGE and stained with mass spectrometry compatible Imperial protein stain (Thermo Fisher Scientific). For the identification of phosphorylation site(s) by PKA, purified Q23-huntingtin (5 µg) was purified in the presence of complete protease and phosphatase inhibitor cocktails (Roche Applied Science) and incubated with or without PRKACA (20 ng). Huntingtin bands (fragments or full-length) of all samples above were excised from gel and processed for mass spectrometry. Excised gel bands were cut into approximately 1 mm^3^ pieces. The samples were reduced with 1 mM DTT for 30 minutes at 60 °C and then alkylated with 5 mM iodoacetamide for 15 minutes in the dark at room temperature. Gel pieces were then subjected to a modified in-gel trypsin digestion procedure (Shevchenko, Wilm, Vorm, & Mann, 1996). LC-MS/MS analysis of the digests was carried out on an LTQ-Orbitrap mass spectrometer (Thermo Finnigan). The eluted peptides were detected, isolated, and fragmented to produce a tandem mass spectrum of specific fragment ions for each peptide. Peptide sequences (and hence protein identity) were determined by matching protein or translated nucleotide databases with the acquired fragmentation pattern by the software program TurboSEQUEST v.27 (Thermo Finnigan) (Eng, McCormack, & Yates, 1994). The modification of 79.9663 mass units to serine, threonine, and tyrosine was included in the database searches to determine phosphopeptides. Each phosphopeptide that was determined by the Sequest program was also manually inspected to ensure confidence. Phosphopeptides obtained were manually aligned to the huntingtin sequence to generate the coverage map.

### Proteome and phosphoproteome profiling with HD neuronal progenitor cells

A panel of twelve HD NPCs (hNPC.01 ∼ 12), each in duplicates were generated from iPSCs following standardized STEMdiff^TM^ neural induction media method (STEMCELL Technologies) except hNPC.02 (neuroal rosettes and FACS sorting (Madison et al., 2015; Sheridan et al., 2011)) and hNPC.04 (Enstem A hNPC purchased from MilliporeSigma) as previously described (Consortium, 2020) and named from the parental iPSC: HD17m.1 (CAG 17/15; NINDS iPSC ID: ND38555), HD17m.8c1 (CAG 17/17), HD19m.4 (CAG 19/16), HD25m.1 (CAG 25/17), HD33i.8 (CAG 33/18; NINDS iPSC ID: ND36997), HD42m.1 (CAG 42/20; NINDS iPSC ID: ND38548), HD45m.2 (CAG 45/15), HD50m.1 (CAG 50/38), HD56m.4 (CAG 56/19), HD60i.4 (CAG 60/18; NINDS iPSC ID: ND36998), HD72m.2 (CAG 72/15), HD180i.7 (CAG 200 ∼240/18; NINDS iPSC ID: ND36999), respectively (Consortium, 2012). NPC cells were cultured on a poly-l-ornithine- and laminin-coated six-well plate (Falcon) at 37°C in neural expansion media (STEMCELL Technologies, 70% DMEM, 30% Hams F12, 1X B27 Supplement, 1% penicillin/ streptomycin, with 20 ng/ml fibroblast growth factor (FGF), 20 ng/ml epidermal growth factor (EGF), and 5 mg/ml heparin freshly added just before use) and washed with PBS and harvested for proteome and phosphoproteome profiling.

Proteome and phosphoproteome profiling of twelve hNPCs, were performed as previously described (Mertins et al., 2018). Briefly, cells were lysed, reduced, alkalated and digested with LysC/Trypsin. Digested peptides were labeled with tandem mass tag (TMT) 10-plex reagent. Three TMT 10-plex experiments were performed each containing 4 of the cell lines in duplicate along with a common reference that was created by pooling equal amounts of all 24 samples. Following successful labeling reactions were quenched and samples for each plex were mixed and desalted. Resulting sample for each plex was first enriched by phosphotyrosine containing peptides using P1000 anti-phosphotyrosine antibody (CST) and enriched fraction was analyzed by LC-MS/MS (Keshishian et al., 2021). Flow through of the enrichment was desalted and fractionated on a 3.5 μm Agilent Zorbax 300 Extend-C18 column (4.6 mm ID x 250 mm length) into 24 fractions. Five percent of each fraction representing total proteome was analyzed by LC-MS/MS (S. Figure 1). Remeining 95% of each of the fraction was enriched by immobilized metal-affinity chromatography (IMAC) for the analyses of phosphoproteome by LC-MS/MS. (S. Table 3). LC-MS/MS analysis of all the samples were performed on QE plus MS system (Thermo Fisher Scientific) as described previously (Mertins et al., 2018).

### Quantitative proteome and phosphoproteome analysis of huntingtin in HD neuronal progenitor cells

All the MS data were searched using Spectrum Mill MS Proteomics Software (Broad Institute) against Uniprot Human database downloaded in October, 2014. Protein and phosphosite level ratios of each sample channel to the common reference was used for further statistical analysis of proteome and phosphoproteome datasets, respectively. A total of 11,998 protein isoforms were found to be detected in all sample replicates. These proteins were used to compare the human NPC samples to one abother. The protein relative abundance levels were quantile normalized between samples using the Bioconductor limma package (3.42.2). After normalization, proteins exhibiting little variantion between samples were removed by filtering away those with a sample-sample varaince of <0.2. This reduced the dataset from 11,998 to 2,694 proteins. Principal components were then calculated from this reduced dataset using R version 3.6.3 and the scatter plot (S. Figure 2) was generated using ggplot2 (3.3.3). The relative adbunace levels of huntingtin phosphopeptides (Figure 2) and peptides (S. Figure 3) were plotted graphically relative to the sequence of the full length huntingtin sequence. The relationship between the relative expression of PRKACA (Uniprot: P17612) and the relative expression of huntingtin (Uniprot: P42858) of tested cell lines was plotted using using R 3.6.3 and ggplot2 3.3.3.

### Kinase panel activity screens with purified huntingtins

A total of 245 Serine/Threonine kinases and 94 Tyrosine kinases were screened to identify kinases targeting purified huntingtins performed by the KinaseFinder Screening service (Reaction Biology). In brief, 5 µg of purified Q23-huntingtin or Q78-huntingtin was incubated with kinases in 50 μl of buffer containing 60 mM HEPES-NaOH, pH 7.5, 3 mM MgCl_2_, 3 mM MnCl_2_, 3 μM Na-orthovanadate, 1.2 mM DTT, 50 μg/ml PEG20000, 1 µM ATP/[γ-^33^P]-ATP (8.24 x 10^05^ cpm per well), protein kinase (1-400 ng/50μl) in 96-well, V-shaped polypropylene microtiter plates at 30°C for 1 h. The reaction was stopped with 20 μl of 10% (v/v) H_3_PO_4_. Of note, we did not use huntingtin purified under conditions that dephosphorylated residue because globally dephosphorylated huntingtin was rapidly insoluble (data not shown). Each sample was transferred into 96-well glass fiber filter plates (MilliporeSigma) which is pre-wetted with 150 mM H_3_PO_4_, followed by incubation at room temperature for 10 min. The plates were washed thrice with 250 μl of 150 mM H_3_PO_4_ and once with 20 μl of 100% ethanol, then dried at 40°C for 30 min. 50 μl of scintillator (CARL ROTH) was added to each well and incorporation of ^33^Pi was measured by a microplate scintillation counter Microbeta (Perkin Elmer). In order to evaluate the results, the background value (C) of the protein was subtracted from raw activity values (A) of each kinase, followed by normalized by the autophosphorylation activity (B) of each kinase which had previously been determined in three independent experiments: Activity ratio of Q23-huntingtin or Q78-huntingtin = (A−C)/B

### *In vitro* kinase assay with recombinant full-length huntingtin and peptides

For radiation detection method, 10 ng of PRKACA (SignalChem) was incubated with 5 µg of huntingtin in a buffer containing 50 mM Tris-HCl pH 7.5, 10 mM MgCl_2_, 0.1 mM EDTA, 2 mM DTT, and 2 µCi ATP/[γ-^32^P]-ATP at 30°C for 1 h. The reaction was stopped by boiling at 95°C for 10 min. Phosphorylated huntingtin was analyzed by SDS-PAGE and visualized by autoradiography with a Typhoon FLA 7000 (GE healthcare).

For immunoblotting method, each 2, 4 µg of GST tagged huntingtin peptides containing S2550 residue (^2546^GRKLSIIRG^2554^) or 5 µg of huntingtin protein was incubated with 1 ng of PRKACA in a buffer containing 50 mM Tris-HCl pH 7.5, 10 mM MgCl_2_, 0.1 mM EDTA, 2 mM DTT, and 200 µM ATP at 30°C for 1 h. Phosphorylated huntingtin peptides or huntingtin are subjected to SDS-PAGE and analyzed by immunoblotting. GST-tagged huntingtin peptide containing S2550 was cloned in pGEX4T1 vector and expressed in *E. coli* BL21 (DE3) cells expression system and purified through glutathione agarose resin. Peptides were eluted from resin in a buffer containing 50 mM Tris-HCl pH 7.5, 100 mM NaCl, 50 mM Glycerol, and 10 mM Glutathione reduced (GSH).

### Human recombinant full-length huntingtin and PRKACA cDNAs in a mammalian expression vector

All recombinant human FLAG tagged huntingtin cDNAs of Q23-huntingtin, Q23-huntingtin carrying S2550A mutation, and Q78-huntingtin used in this study were cloned in a modified pcDNA3 vector. The original polyclonal region of pcDNA3 vector (Invitrogen) was swapped with the modified polyclonal region containing 1X FLAG, 6X histidine tag, TEV protease recognition site, and several restriction enzyme sites, HindIII, BamHI, XhoI, SacII, and ApaI. Human FLAG tagged Q23-huntingtin and Q78-huntingtin cDNAs from previously reported human huntingtin pALHDQ23 insect cell expression vector (Shin et al., 2018) was inserted into between BamHI and SacII. To generate Q23-huntingtin S2550A cDNA, site-directed mutagenesis was carried out using Quick Change site-directed mutagenesis kit (Agilent) with pcDNA3 FLAG Q23-huntingtin as template and mutagenesis primer, 5’-GGGAGGAAGCTGGCGATTATCAGAGGG-3’ according to the protocol from the manufacture. The recombinant human PRKACA cDNA was obtained from Addgene (Plasmid #23495) and cloned into between BamHI and KpnI in pcDNA5 vector (Invitrogen). All final clones were verified using full DNA sequence analysis.

### Cell culture, transfection and treatments

*HTT* null HEK 293T, which was previously generated by using CRSPR/Cas9 to remove the first exon and upstream promoter region of *HTT* (R. Jung et al., 2021) and parental HEK 293T cell lines were maintained in Dulbecco’s Modified Eagle’s Medium (Invitrogen) supplemented with 1% Penicillin-Streptomycin (Gibco) and 10% FBS (MilliporeSigma) at 37°C in a humidified 5% CO_2_ atmosphere. HD60i.4 NPCs (CAG 60/18) were cultured in neural expansion media as described above.

All recombinant human huntingtin and PRKACA cDNA plasmids were transfected into HEK 293T cells or *HTT* null HEK 293T cells with Lipofectamine 3000 (Invitrogen) following manufacturer’s instructions. 30 µM of cycloheximide (MilliporeSigma) was treated to cells expressing huntingtin without or with PRKACA at post 24 hours transfection. 300 µM of 8-Br-cAMP (Santa Cruz Biotechnology) was treated to hNPCs for 0, 4, 8, 24 hours to trigger PKA activity.

### Immunoblotting

Each sample was loaded on NuPAGE™ 4-12% Bis-Tris Protein Gels (Invitrogen) and separately by applying 120 V for 120 min except Figure 6C where the gel ran at 120 V for 300 min to separate Q20-huntingtin and Q62-huntingtin. Proteins were transferred to PVDF or nitrocellulose membrane from gels on demand. The membrane was blocked in a Tris-buffered saline buffer containing 0.1% tween 20 (TBS-T) with 5% skim milk for 1 hour or 5% BSA (for phosphor antibodies) for overnight. Each primary antibody (see below) was diluted in blocking buffer according to the manufacturer’s instructions and incubated with membrane at 4°C overnight. After washing membrane three times using TBS-T buffer, each secondary antibody was incubated with membrane in a TBS-T buffer at room temperature for 1 h, and then washed three times with TBS-T buffer. For signal detection, Western lightning ECL pro (PerkinElmer) was used to film development.

Primary antibodies used in this research are as follows: mouse monoclonal anti-huntingtin antibody MAB2166 (MilliporeSigma), MAB2168 (MilliporeSigma), 1F8 (White et al., 1997), rabbit polyclonal anti-phospho Serine 2550 antibody (abHTT-pS2550) (T. Jung et al., 2020), rabbit polyclonal anti-GAPDH antibody (Santa Cruz Biotechnology), mouse monoclonal anti-α-tubulin antibody (Cell Signaling Technology), mouse monoclonal anti-PRKACA antibody (Santa Cruz Biotechnology), mouse monoclonal anti-CREB antibody (Cell Signaling Technology), rabbit monoclonal anti-phospho CREB (S133) antibody (Cell Signaling Technology).

### Parallel reaction monitoring (PRM) MS analysis to quantify S2550 phosphopeptide

To enrich endogenous huntingtin in cells in the absence or presence of exogenous PRKACA expression (24 hour), cell were harvested and lysed in a lysis buffer containing 0.2% n-dodecyl-β-D-maltoside, 5 mM Mg(OAC)_2_, 70 mM KOAc, 50 mM HEPES pH 7.5, and cOmplete™ protease inhibitor cocktail tablet (Roche Applied Science) on a rotor at 4°C for 30 minutes. The lysed cells were centrifuged for 15 minutes at 13000 rpm and then the supernatants were incubated mouse monoclonal anti-huntingtin antibody MAB2166 (MilliporeSigma) conjugated Protein G Agarose (Roche) at 4°C for 2 hours. The precipitants were washed with the lysis buffer thrice. The bound samples were boiled to elute from resin and used for immunoblotting and tandem mass spectrometric analysis. To quantify S2550 phosphopetides, PRM analyses were performed on a Q-Exactive mass spectrometer equipped with an Easy nLC-1000 (Thermo Fisher Scientific) at Quantitative Proteomics Resource Core at University of Pennsylvania Medicine. The peptides samples were separated using a linear gradient of 2% -35% solvent B (0.1% formic acid in acetonitrile) at a flow rate of 300 nL min-1 over 40 minutes, followed by an increase to 90% B over 4 minutes and held at 90% B for 6 min before returning to initial conditions of 2% B. For peptide ionization, 2000 V was applied and a 250 °C capillary temperature was used. All samples were analyzed using a multiplexed PRM method based on an unscheduled inclusion list containing the target precursor ions and heavy isotope-labeled peptides. The full scan event was collected using an m/z 380–1500 mass selection, an Orbitrap resolution of 70K (at m/z 200), a target automatic gain control (AGC) value of 1 × 106, and maximum injection time of 54 milliseconds. The PRM scan events used an Orbitrap resolution of 17,500, an AGC value of 1 × 106, and a maximum fill time of 64 milliseconds with an isolation width of 2 m/z. Fragmentation was performed with a normalized collision energy of 27 and MS/MS scans were acquired with a starting mass of m/z 140. 100 fmol of heavy isotope-labeled peptides (KLSIIR*) was spiked into each sample. PRM data analysis was performed using Skyline software (MacLean et al., 2010).

### Lentiviral transduction

MISSION® shRNA PKA (shPKA, MilliporeSigma, SHCLNG-NM_002730, TRCN0000367487) was used to knock down cellular PRKACA level. The sequences for PRKACA shRNA was 5’- CCGGGATAATCAGAGGGACAGAAACCTCGAGGTTTCTGTCCCTCTGATTATCTTTTTG-3’. Lentiviral particles encoding shPKA were transduced into HEK 293T cells at various range of a multiplicity of infection (MOI) in the presence of 8 µg/ml of polybrene to find optimal degree of PKA knock down. Cells were selected with puromycin by examining viability every 2 days for 14 days to genearate HEK 293T cells stably expressing shRKA RNA. To evaluate the reduced RNA expression level of PRKACA in selected stable cells, total RNA was prepared by RNeasy Plus kit (Qiagen) and cDNA synthesis was performed using SuperScript IV First-Strand Synthesis system (Thermo Scientific). RRKACA RNA expression level was measured by Quantitative real-time PCR (qRT-PCR) using LightCycler 480 SYBR Green I Master kit on Roche LightCycler 480 instrument. In order to detect protein expression level of PRKACA and huntingtin in selected stable cells, western blot was used as described above in methods.

## ACKNOWLEDGEMENTS

We thank the members of the Song and Seong laboratories for suggestions and discussions. We are grateful to Dr. Ross Tomaino (The Taplin Mass Spectrometry Facility) for excellent technical support and also thank Hyungjoo Lee at Quantitative Proteomics Resource Core of School of Medicine at the University of Pennsylvania. This work was supported by a Global Research Laboratory grant [NRF-2016K1A1A2912057 to ISS and JS] and [NRF-2020R1A2B5B03001517 to JS] from National Foundation of Korea and CHDI Foundation [MEM and ISS].

## AUTHOR CONTRIBUTIONS

YL, DB, HK, KH, RV, RSA, HS, HK, JL designed and conducted experiments and performed data analysis. JS, ISS, SC, RL, SK, MEM contributed to the design of the study and YL, MEM, JS and ISS were involved in writing the manuscript, which was reviewed by all the authors.

## COMPETING FINANCIAL INTERESTS

S.A.C. is a member of the scientific advisory boards of Kymera, PTM BioLabs, Seer and PrognomIQ. J.S. is a co-founder of PCG-Biotech, Ltd. The remaining authors declare no competing financial interests.

## Supplementary Figure Legends

**S. Figure 1.**
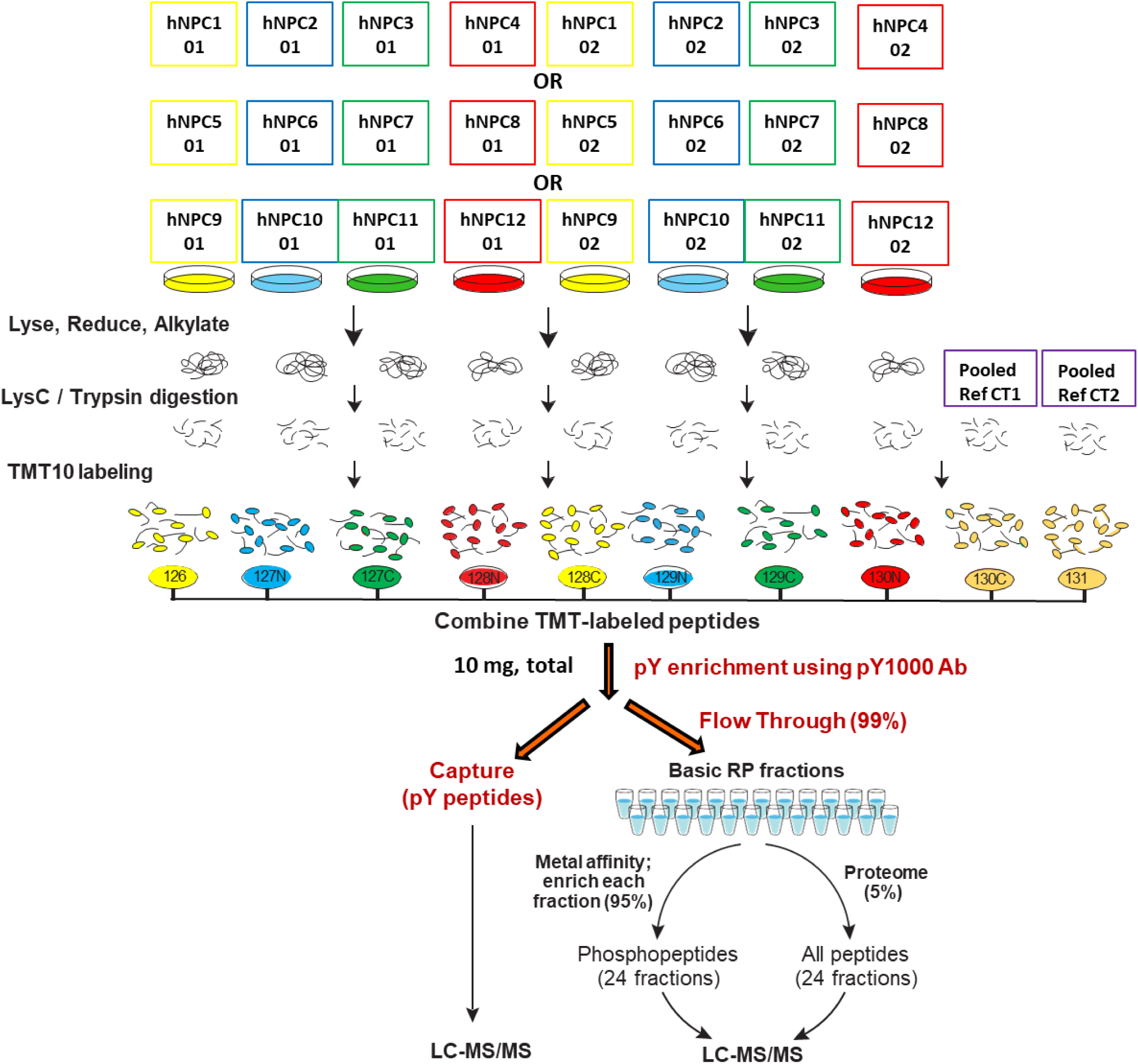
Workflow and experimental design of proteome and phosphoproteome of a panel of 12 HD hNPCs using TMT. The top lines of the flow diagram show the sample pooling scheme; using TMT 10-plex reagents, duplicates of four hNPCs (hNPC.01 ∼ hNPC.04) plus duplicates of pooled reference control that was mixed with each same portion lysates of all tested hNPC lysastes were performed, along with two more sets (hNPC.05 ∼ hNPC.08 and hNPC.09 ∼ hNPC.012) that were subjected into the same proteomic and phosphoproteomic approaches. The bottom portion outlines steps for cell lysates with phosphotyrosine antibody enrichment, fractionation, metal affinity for phosphoprotein enrichment and quantitagive MS analysis.

**S. Figure 2.**
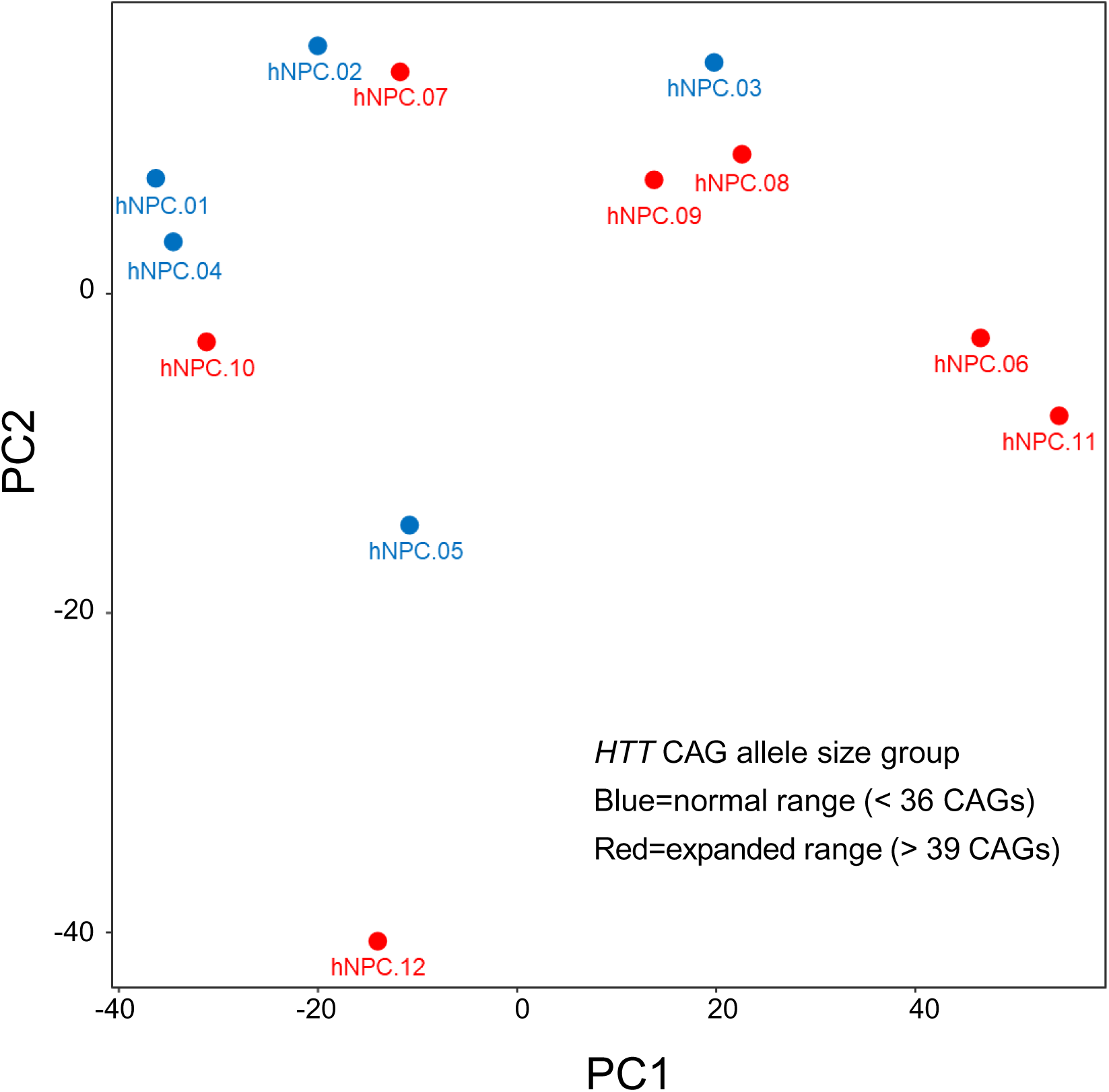
Principal component protein abundance across the 12 sample hNPC dataset. Principal components were calculated and plotted from protein abundance values after removing proteins exhibiting relatively invariant (variance < 0.2) abundance, revealed that hNPC.12 was an outlier compared to the other members of the panel.

**S. Figure 3.**
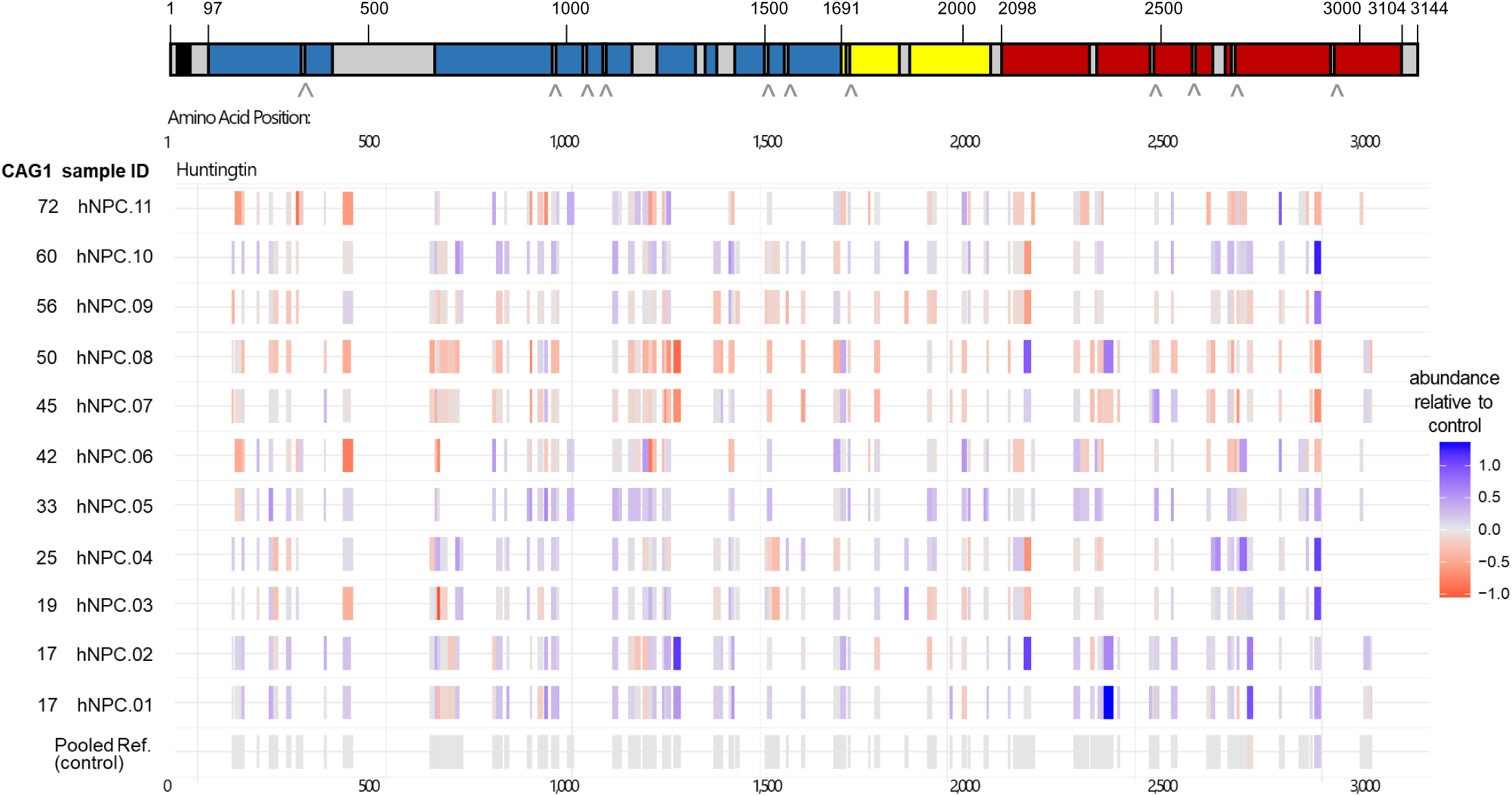
Location and relative abundance of huntingtin peptides identified in eleven hNPCs. The location and relative abundance of the peptides identified from eleven hNPCs under a schematic of full length huntingtin, with its N-HEAT (blue), Bridge (yellow) and C-HEAT (red) domains, with structurally unresolved regions denoted in grey, for 11 members of the hNPC panel, ranked by size of longer CAG repeat allele (left column), with abundance of each peptide relative to the pooled reference control denoted by the heat-scale.

**S. Figure 4.**
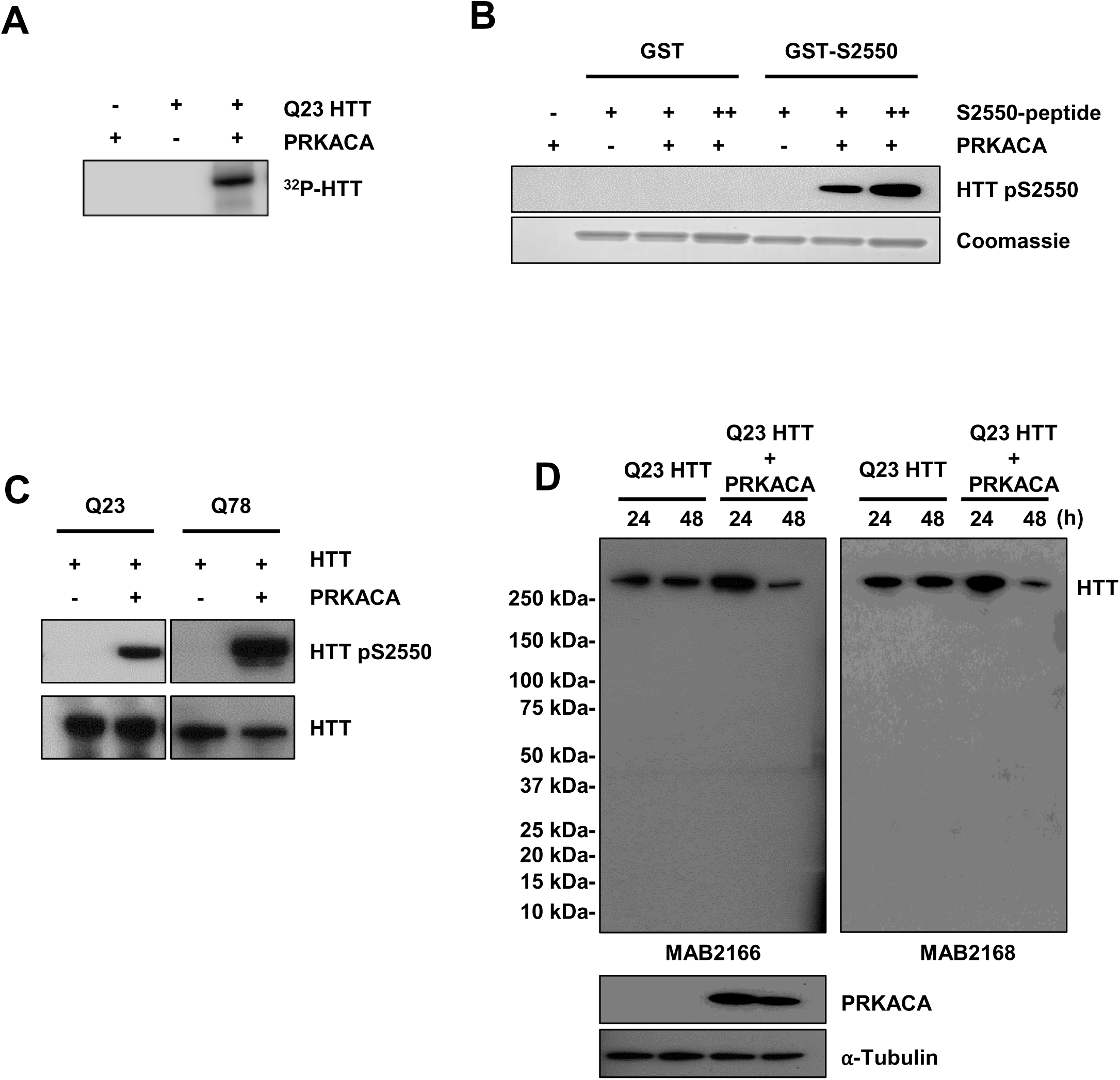
Validation of huntingtin Ser2550 as target site by PKA. A. An autoradiogram of SDS-PAGE showing a band of phosphorylated huntingtin (HTT) labelled with [γ-^32^P] ATP only in PRKACA **B.** The immunoblot shows a band of phosphorylated Ser2550-GST-tag-huntingtin peptide containing Ser2550 residue incubated with PRKACA but not in GST-tag only. Bottom panel shows the equal amounts of each peptide visualized with Coomassie Blue staining. **C.** An immunoblot showing bands of pSer2550-purified Q23-huntingtin and pSer2550Q78-huntingtin detected with abHTT-pS2550 antibody only after *in vitro* co-incubation with PRKACA (top), with bands of total huntngtin detected by MAB2166 (bottom). **D.** Immunoblots showing detection of bands of overexpressed huntingtin without or with PRKACA overexpression in *HTT* null HEK 293T cells for 24 hours or 48 hours, detected with two anti-huntingtin reagents; MAB2166 epitope a.a. 181-810 of huntingtin (left), with PRKACA level detected with anti-PRKACA reagent and α-tubulin as loading control with anti-α-tubulin reagent.in the panel below, and MAB2168 epitope region: a.a. 2146-2541 (right). Note that no shorter products were detected by either reagent.

**S. Table 1:** Unresolved huntingtin regions in the human hutingtin/HAP40 cryo-EM structure (PDB: 6EZ8)

| Unresolved huntingtin region (amino acid number: start-end)* | Renumbered for Q23-huntingtin | Length of unresolved region (# residues) | Domain location |
| --- | --- | --- | --- |
| 1–90 | 1–96 | 96 | N-HEAT |
| 323–342 | 329–348 | 20 | N-HEAT |
| 403–660 | 409–666 | 258 | N-HEAT |
| 960–977 | 966–983 | 18 | N-HEAT |
| 1,049–1,057 | 1055–1063 | 9 | N-HEAT |
| 1,103–1,120 | 1109–1126 | 18 | N-HEAT |
| 1,158–1,222 | 1164–1228 | 65 | N-HEAT |
| 1,319–1,347 | 1325–1353 | 29 | N-HEAT |
| 1,372–1,418 | 1378–1424 | 47 | N-HEAT |
| 1,504–1,510 | 1510–1516 | 7 | N-HEAT |
| 1,549–1,556 | 1555–1562 | 8 | N-HEAT |
| 1,714–1,728 | 1720–1734 | 15 | Bridge |
| 1,855–1,881 | 1861–1887 | 27 | Bridge |
| 2,063–2,091 | 2069–2097 | 29 | Bridge |
| 2,325–2,347 | 2331–2353 | 23 | C-HEAT |
| 2,472–2,490 | 2478–2496 | 19 | C-HEAT |
| 2,580–2,582 | 2586–2588 | 3 | C-HEAT |
| 2,627–2,660 | 2633–2666 | 34 | C-HEAT |
| 2,681–2,687 | 2687–2693 | 7 | C-HEAT |
| 2,926–2,944 | 2932–2950 | 19 | C-HEAT |
| 3,099–3,138 | 3105–3144 | 40 | C-HEAT |
| **Total unresolved region residues:** |  | 791 |  |
| *Numbering for 17-polyglutamine huntingtin |  |  |  |

**S. Table 2:**
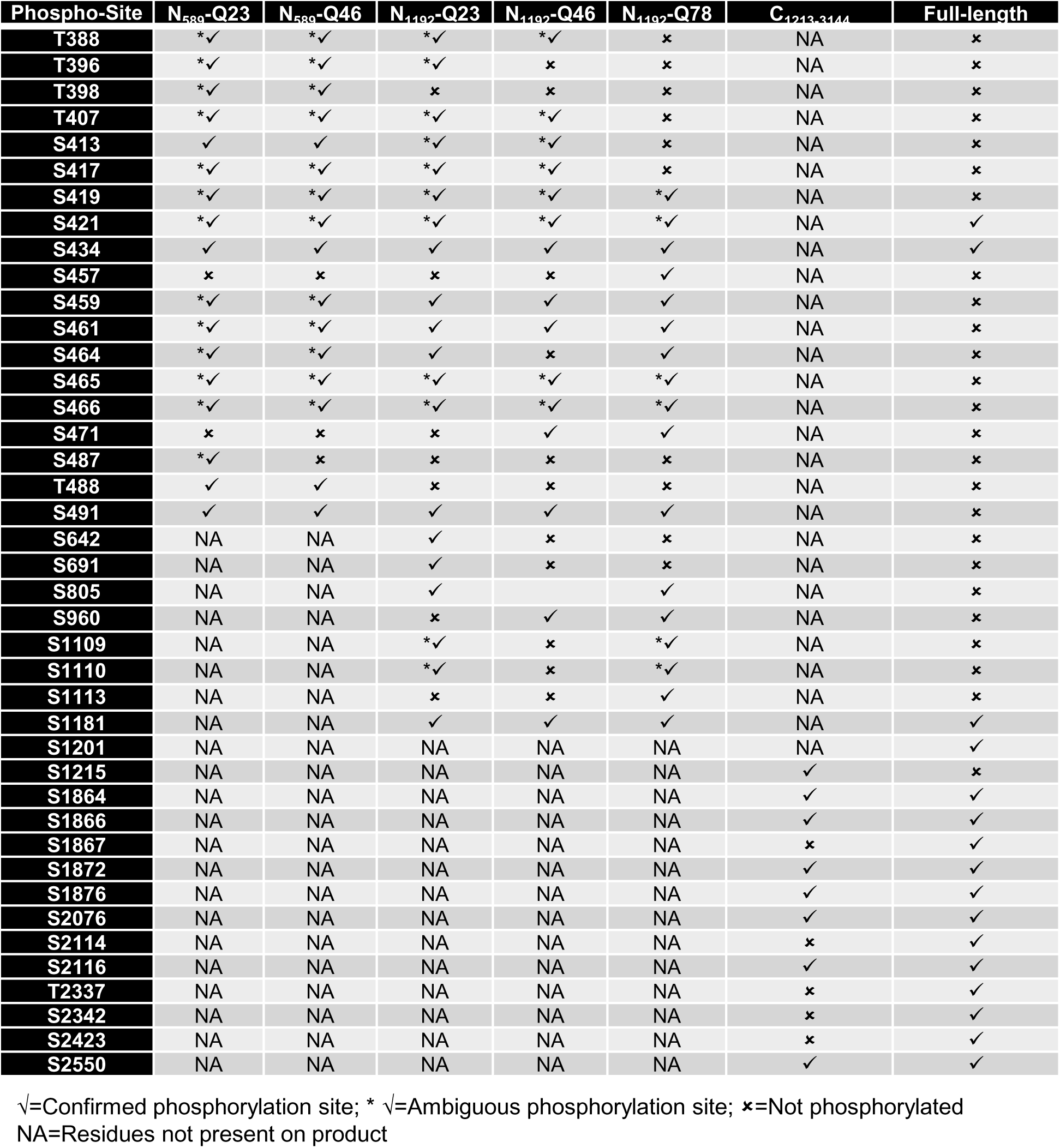
Comparison of phosphorylation sites.

**Three Supplemental tables below included as three excel files**

**S. Table 3. Huntingtin peptides and phosphopeptides detected from HD hNPC panel**

**S. Table 4. Huntingtin PhosphoSite mass spec all studies.**

**S. Table 5. Q23-huntingtin Q78-huntingtin kinase activity data**

